# Three-Year Longitudinal Analysis of Splicing Changes in Myotonic Dystrophy Type 1

**DOI:** 10.64898/2026.09.25.754454

**Authors:** Cécilia Légaré, Lori Planco, Robert Merritt, John Ripollone, Sarah Conner, Laura M. Desrochers, Fares Nigim, Marie-Pier Roussel, Cynthia Gagnon, John Douglas Cleary, J. Andrew Berglund, Elise Duchesne

## Abstract

Myotonic dystrophy type 1 (DM1) is caused by a CTG expansion in the 3′ untranslated region of the *DMPK* gene that results in the expression of expanded CUG RNA which trap splicing factors resulting in widespread alternative splicing dysregulation. A panel of these splicing events, or splicing index (SI), is currently used to measure splicing dysregulation in DM1 patients. Despite significant interest in splicing, little is known about its progression over time in DM1. The objective of this study was to evaluate the changes in SI over 3 years in a well-phenotyped cohort of DM1 participants and identify factors contributing to these changes. RNA was extracted from *vastus lateralis* (VL) biopsies of 20 DM1 participants taken 3 years apart and the SI score was measured using targeted RNA sequencing. Baseline SI scores correlated with percent predicted muscle strength of the ankle dorsiflexors, knee extensors and hand grip. On average, SI scores worsened over 3 years, but the progression varied markedly among participants. Adult/juvenile phenotypes and lower baseline SI scores were among the factors associated with higher progression of SI scores over time. These findings emphasize the importance of integrating clinical phenotype and baseline molecular severity when interpreting longitudinal SI change, particularly in the context of clinical trials.

## Introduction

A CTG repeat expansion in the 3′ untranslated region of the *dystrophia myotonica protein kinase* (*DMPK*) gene is responsible for myotonic dystrophy type 1 (DM1)(1–3). While the global prevalence of DM1 is 1 in 8000, some regions can be as high as 1 in 2100 (New York state, USA)(4) or 1 in 630 (Saguenay–Lac-St-Jean region in Quebec, Canada)(5). DM1 is a complex, multisystemic disease affecting nearly all organ systems, including muscular, central nervous, ocular, gastrointestinal, endocrine, cardiovascular and respiratory systems(6). This highly variable and heterogeneous disease is currently divided into five clinical phenotypes based on the number of CTG repeats and age at onset: congenital, childhood, juvenile, adult, and late-onset(7).

Expression of the *DMPK* CUG expansion RNA traps splicing regulators such as muscleblind-like (MBNL) proteins, resulting in global splicing alterations(8–11). While there are currently no approved disease-modifying treatments for DM1, several ongoing clinical trials (NCT06185764, NCT05027269, NCT06667453) target the toxic DMPK RNA, with correction of RNA splicing abnormalities being evaluated as a pharmacodynamic biomarker of target engagement and treatment response(12). To assist in assessing splicing, a splicing index score (SI) was developed from *tibialis anterior* (TA) muscle samples using 22 individual splicing events that are associated with either DM1 pathology and/or clinical disease severity(13). Based on an analysis of 58 DM1, 22 unaffected and 11 disease controls, Provenzano *et al*, separated the SI score into three categories based on splicing dysregulation: mild (SI ≤0.4), moderate (SI: 0.4-0.75) and severe (SI ≥0.75)(14). They found that over 3 months, the SI score increased, with participants in the moderate group showing the biggest change. While this study established the SI as a promising biomarker, little is known about the progression of SI over longer periods of time or in muscles other than the TA.

Our group previously demonstrated histopathological changes and clinical decline over 3 years in individuals with DM1, including a significant reduction in muscle strength in hand grip (−17.4%), pinch grip (−30.6%), hip abductors (−17.4%), knee extensors (−8.5%) and ankle dorsiflexors (−8.4%)(15, 16). While informative, neither these studies nor others have examined the longitudinal progression of splicing alterations or the factors that may influence these changes over long periods of time. Therefore, the objectives of this study were to use *vastus lateralis* (VL) muscle biopsies from a well characterized DM1 cohort to: 1) compare to previously reported associations between SI scores measured in TA muscle biopsies and clinical outcomes; 2) assess the progression of splicing alterations over 3 years; and 3) identify factors contributing to change in these splicing alterations. Here we show that baseline SI scores from VL correlated with percent predicted muscle strength in the ankle dorsiflexors, knee extensors and hand grip. On average, these SI scores worsened over 3 years, with progression varying markedly among participants. Adult/juvenile phenotype and lower baseline SI score were among the factors associated with higher progression of SI scores over time. These findings highlight that interpretation of longitudinal SI change requires integration of clinical phenotypes and baseline molecular severity.

## Results

### Associations between splicing index and clinical measures at baseline

Participants’ characteristics were collected at start of the study (baseline) for the cohort of 20 DM1 participants, which consisted of 11 women and 9 men with an average age of 51.2 years (**Table 1**). Clinical phenotypes were reviewed and assigned by a neuromuscular neurologist, with the single participant with the infantile phenotype integrated in the juvenile group for analysis. In our cohort, more participants have juvenile (35%) and adult (45%) phenotypes than the late-onset phenotype (20%). As expected, participants with the juvenile and adult phenotypes have lower muscle strength (expressed as percent predicted), and higher CTG repeat length and SI score than those with the late-onset phenotype.

**Table 1:** Participants characteristics at baseline.

| Participant's characteristics | Overall | By sex |  | By phenotype |  |  | By repeat length |  |  |
| --- | --- | --- | --- | --- | --- | --- | --- | --- | --- |
|  |  | Women | Men | Juvenile | Adult | Late | <300 | 300-800 | >800 |
| n (%) | 20 | 11 (55) | 9 (45) | 7 (35) | 9 (45) | 4 (20) | 6 (30) | 10 (50) | 4 (20) |
| Age, mean (SD) | 51.2 (11.0) | 49.3 (12.0) | 53.6 (9.7) | 42.0 (12.5) | 54.4 (5.6) | 60.0 (5.4) | 57.8 (5.4) | 48.4 (12.1) | 48.3 (12.1) |
| [min-max] | [27-65] | [27-65] | [32-64] | [27-57] | [45-62] | [54-65] | [52-65] | [27-62] | [32-60] |
| Repeat length, mean (SD) | 538.3 (323.9) | 658.0 (303.0) | 391.9 (300.9) | 751.4 (200.2) | 571.1 (279.6) | 91.3 (6.8) | 150.2 (91.9) | 583.1 (110.4) | 1008 (106.3) |
| [min-max] | [85-1097] | [101-1097] | [85-954] | [570-1097] | [254-1097] | [85-101] | [85-282] | [349-748] | [885-1097] |
| [n women, n men] | [11,9] |  |  | [6,1] | [4,5] | [1,3] | [2,4] | [6,4] | [3,1] |
| [n juvenile, n adult, n late] | [7,9,4] | [6,4,1] | [1,5,3] |  |  |  | [0,2,4] | [5,5,0] | [2,2,0] |
| Knee extensors strength (% predicted), mean (SD) | 51.7 (22.4) | 40.8 (15.9) | 64.9 (22.8) | 38.5 (9.9) | 48.4 (19.3) | 82.1 (18.2) | 70.6 (23.0) | 49.6 (15.8) | 28.6 (10.8) |
| [min-max] | [19.8-109.3] | [19.8-73.6] | [33.3-109.3] | [22.8-49.2] | [19.8-73.2] | [71.2-109.3] | [42.3-109.3] | [31.2-73.2] | [19.8-44.1] |
| Ankle dorsiflexors strength (% predicted), mean (SD) | 55.4 (22.4) | 48.8 (16.1) | 62.7 (26.8) | 45.6 (13.4) | 46.6 (12.3) | 89.8 (17.3) | 76.5 (24.8) | 43.5 (13.2) | 52.7 (11.5) |
| [min-max] | [28.3-109.8] | [28.3-73.4] | [34.7-109.8] | [31.3-66.9] | [28.3-65.2] | [73.4-109.8] | [44.4-109.8] | [28.3-66.9] | [42.4-65.2] |
| Grip strength (% predicted), mean (SD) | 46.5 (28.0) | 44.3 (22.5) | 49.2 (34.9) | 32.8 (17.7) | 36.2 (13.4) | 93.7 (11.5) | 76.5 (28.4) | 34.0 (15.3) | 32.8 (18.8) |
| [min-max] | [9.7-104] | [9.8-94.2] | [12.5-104.0] | [9.8-59.6] | [12.5-52.0] | [77.5-104.0] | [35.4-104.0] | [12.5-59.6] | [9.8-52.0] |
| SI score, mean (SD) | 0.3 (0.3) | 0.4 (0.3) | 0.2 (0.3) | 0.4 (0.2) | 0.4 (0.4) | 0.02 (0.1) | 0.06 (0.1) | 0.4 (0.3) | 0.6 (0.3) |
| [min-max] | [-0.04-0.98] | [0.04-0.98] | [-0.04-0.77] | [0.2-0.7] | [0.04-0.98] | [-0.04-0.2] | [-0.04-0.2] | [0.07-0.98] | [0.23-0.89] |

Abbreviations: %: percentage, SD: standard deviation, min: minimum, max: maximum, SI: splicing index.

As SI scores have been shown to correlate with clinical measures in TA muscle(14), we first examined the association between VL baseline SI score and diverse clinical outcomes shown to be valid and reliable in the DM1 population (**Figure 1**). The VL SI score in our cohort is strongly negatively correlated with baseline percent predicted ankle dorsiflexors muscle strength (r_s_= - 0.533, p = 0.019, **Figure 1A**), percent predicted knee extensors muscle strength (r_s_= −0.684, p = 0.001, **Figure 1B**) and percent predicted grip strength (r_s_= −0.457, p = 0.043, **Figure 1C**), but not significantly with 10-meter walk test (10mWT) maximum speed (r_s_= −0.165, p = 0.486, **Figure 1D**). To reflect whole-body impairments (lower limb distal and proximal muscles and upper limb muscles), we created composite scores that combine muscle strength of ankle dorsiflexors, knee extensors/or and grip expressed as percent predicted. These composite scores showed higher correlations with the SI score (**Figure 1E-H**), with the combination of ankle dorsiflexors and knee extensors showing the best correlation (r_s_= −0.705, p = <0.001, **Figure 1E**). The same composite score approach was applied in a secondary analysis of the TA data from Provenzano *et al*, which demonstrated that this composite approach produces higher correlations with SI score than individual muscle strength measures (**Supplemental Figure 1 A-D**).

**Figure 1:**
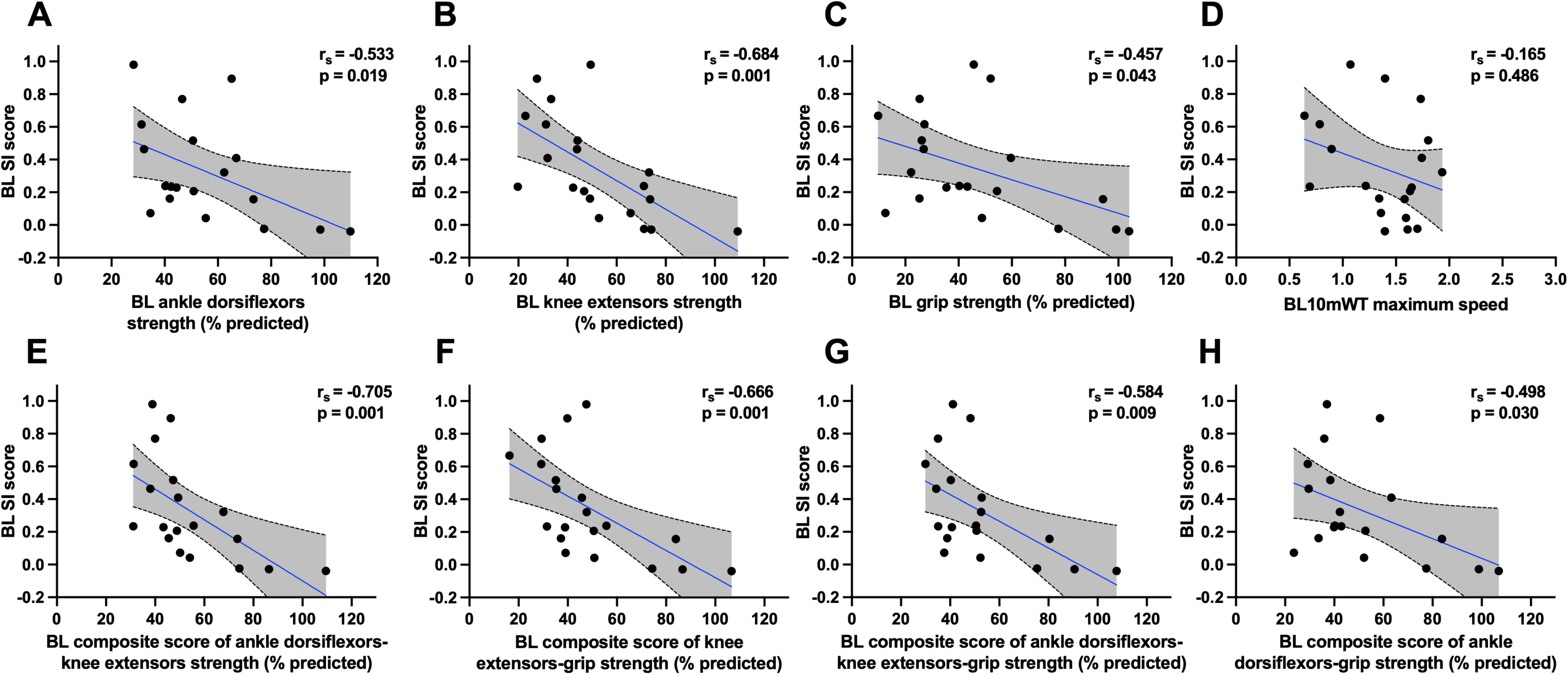
Spearman correlations of splicing index and clinical values at baseline. **A.** Correlation of splicing index and ankle dorsiflexor strength. **B.** Correlation of splicing index and knee extensors strength. **C.** Correlation of splicing index and grip strength. **D.** Correlation of splicing index and 10-meter Walk Test maximum speed. **E.** Correlation of splicing index and composite score average of percent predicted quantitative muscle testing of ankle dorsiflexors and knee extensors strength. **F.** Correlation of splicing index and composite score average of percent predicted quantitative muscle testing of knee extensors and grip strength **G.** Correlation of splicing index and composite score average of percent predicted quantitative muscle testing of ankle dorsiflexors, knee extensors and grip strength. **H.** Correlation of splicing index and composite score average of percent predicted quantitative muscle testing of ankle dorsiflexors and grip strength. Abbreviations: BL: baseline, SI: splicing index, %: percent, 10mWT: 10-meter Walk Test.

### Splicing progression over 3 years

To understand longitudinal changes in splicing and clinical progression, we next compared SI scores over a 3-year time from baseline (**Table 2**). On average, the median SI score significantly increase (worsened) from 0.24 to 0.44 (p = 0.002) over 3 years for all participants (**Supplemental Table 2**), but the progression was highly variable from one participant to another (n = 20, median [Q1, Q3] of change in SI score = 0.088 [0.013, 0.240], **Supplemental Table 2**). To better understand this variability, we conducted a subgroup analysis based on the baseline SI score range (mild: ≤0.4, moderate: 0.4-0.75, severe: ≥0.75) as per Provenzano *et al*. Over three years, participants in the mild SI group had statistically significant increases in SI score, but presented the largest variation with some participants showing no change with others showing the highest changes (n = 12, median [Q1, Q3] = 0.16 [0.025, 0.28], p-value = 0.005). The moderate group had a non-statistically significant trend towards increases in SI score (n=5, median [Q1, Q3] = 0.08 [−0.01,0.3], p-value = 0.188), while the severe SI group had little to no change (n = 3, median [Q1, Q3] = −0.01 [−0.07,0.09], p-value = 0.999) (**Figure 2A**). We also examined each participant based on their determined phenotype (juvenile, adult & late) and interestingly observed that participants presenting with late-onset phenotype (#4, 5, 6 & 20) in the mild group showed little or no SI score progression compared to other participants in the group (**Figure 2A**). We next examined participants based on their CTG repeat length (<300, 300-800, >800) but did not observe a noticeable association with changes in SI score. Overall, participants with a mild SI score at baseline (SI ≤0.4) presented the greatest median changes in SI score over the 3-year period, except participants with the late phenotype that presented the lowest change.

**Figure 2:**
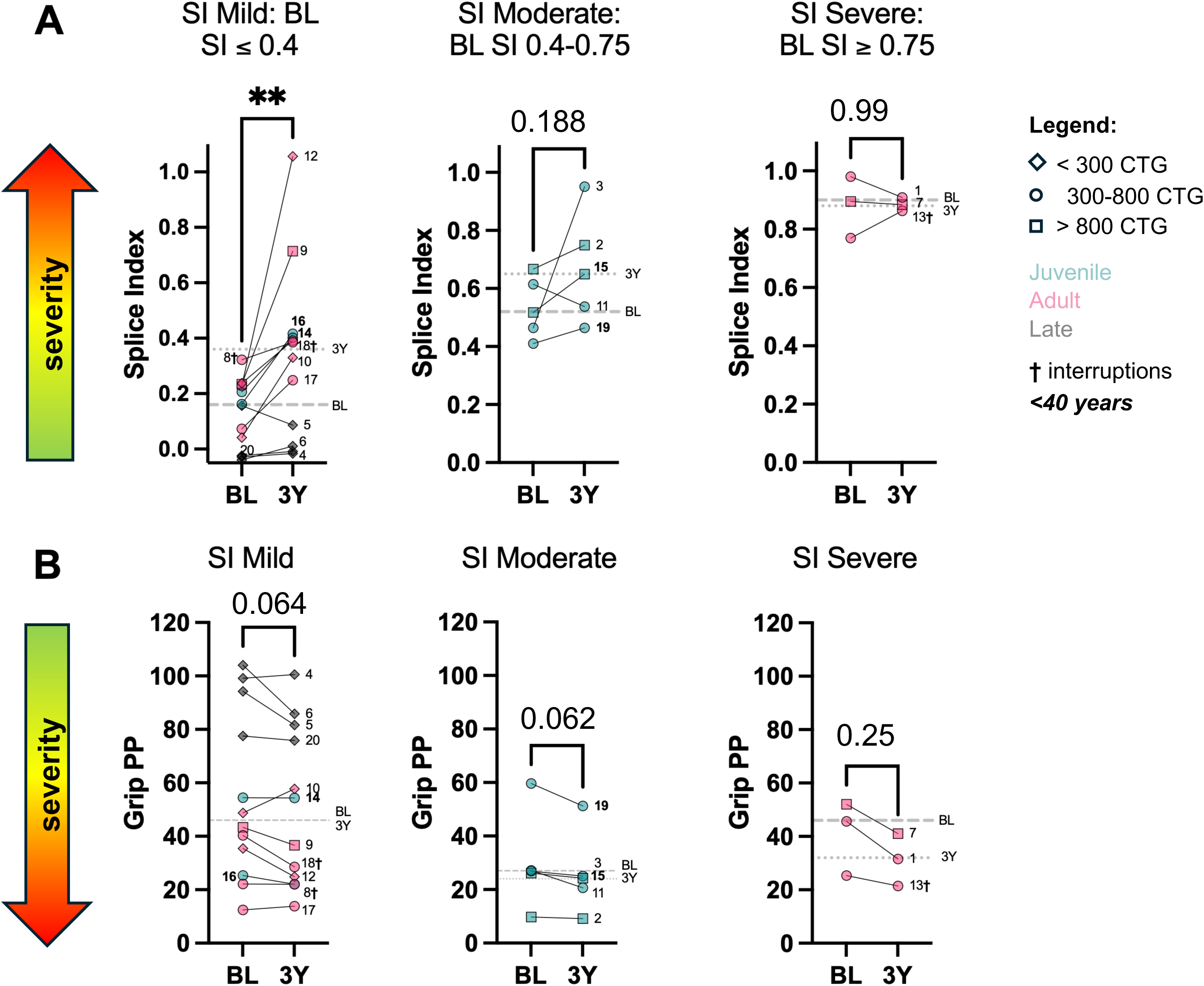
Splice index score and grip strength changes over 3 years. **A.** SI score changes over 3 years in participants. Shapes represent their number of CTG repeats <300 (diamond), 300-800 (circle), >800 (square) while colors represent their phenotype juvenile (green), adult (pink), late (grey), participants with interruption of their repeats have cross next to their ID in their labels and ***bold in italic*** ID labels are participants <40 years old. Median are presented as grey lines large dash= baseline, smaller dash=3 years. **B.** Grip strength changes over 3 years. Shapes represent their number of CTG repeats <300 (diamond), 300-800 (circle), >800 (square) while colors represent their phenotype juvenile (green), adult (pink), late (grey), participants with interruption of their repeats have cross next to their ID in their labels and ***bold in italic*** ID labels are participants <40 years old. Median are presented as grey lines large dash= baseline, smaller dash=3 years. Abbreviations: SI: splice index, BL: baseline, 3Y: 3 years, grip pp: grip percent predicted quantitative muscle testing strength, CTG: number of CTG repeats. **p<0.01 Wilcoxon test.

**Table 2:** Changes in clinical values and SI score over 3 years.

|  | All participants (n=20) |  |  | All participants with SI> 0 at BL (n=17) |  |  |
| --- | --- | --- | --- | --- | --- | --- |
| All participants | Baseline | 3 years | P-value | Baseline | 3 years | P-value |
| SI score, median (Q1, Q3) | 0.2 (0.1, 0.6) | 0.4 (0.3, 0.8) | 0.002 | 0.3 (0.2, 0.6) | 0.5 (0.4-0.9) | 0.002 |
| [min-max] | [-0.04-0.98] | [-0.02-1.06] |  | [0.04-0.98] | [0.09-1.06] |  |
| Knee extensors strength (Nm), median (Q1, Q3) | 57.3 (46.1, 97.0) | 50.3 (30.6, 79.2) | <0.001 | 53.8 (36.0, 79.2) | 48.0 (27.8, 76.1) | 0.003 |
| [min-max] | [16.2-176.6] | [15.0-146.4] |  | [16.2-127.4] | [15.0-117.0] |  |
| Ankle dorsiflexors strength (Nm), median (Q1, Q3) | 12.0 (8.3, 16.0) | 10.6 (5.6, 16.0) | 0.060 | 10.6 (7.2, 13.2) | 9.9 (5.4, 12.1) | 0.160 |
| [min-max] | [5.5-31.6] | [1.7-29.1] |  | [5.5-17.9] | [1.7-20.8] |  |
| Grip strength (kg), median (Q1, Q3) | 13.6 (8.3, 18.5) | 10.6 (6.8, 16.9) | <0.001 | 10.7 (7.7, 14.9) | 8.9 (6.6, 13.3) | <0.001 |
| [min-max] | [2.7-50.8] | [2.2-37.9] |  | [2.7-19.3] | [2.2-17.3] |  |

Abbreviations: SI: splicing index, Q1: quartile 1, Q3: quartile 3, min: minimum, max: maximum, BL: baseline.

### Factors influencing changes in SI score over 3 years

To identify factors contributing to the worsening in SI score over 3 years, multiple linear regression models were created (**Table 3 & Supplemental Table 1**), which incorporated phenotype, progression of SI score, and baseline SI score. Additional variables were also examined if they showed a Spearman correlation coefficient of 0.2 or more with delta SI (**Supplemental Figure 2**). Adult and juvenile onset phenotypes (Standardized β = −0.683, p = 0.008) and lower SI score at baseline (Standardized β= - 0.607, p = 0.016) were both associated with greater worsening in the SI score, these two variables explaining 31% of the variance in change in SI score (**Table 3 model A**). The identification of baseline SI score and clinical phenotype as factors influencing SI score progression over time prompted a more detailed examination of the relationship between these variables (**Figure 3**). We observed that most late-onset participants (3 out of 4) had no evidence of splicing dysregulation at baseline (i.e. baseline SI scores below 0) which could weaken the overall association between baseline SI score and SI progression (r = −0.182). Exclusion of these participants (4, 6 & 20) substantially strengthened the correlation between baseline SI score and changes in SI score over three years (**Figure 3B**; r = −0.529, p = 0.031). Additionally, according to the linear regression models, participants with higher baseline SI scores (#1, 7 & 13) are associated with smaller changes in SI over time, suggesting a ceiling effect in the progression of mis-splicing.

**Figure 3:**
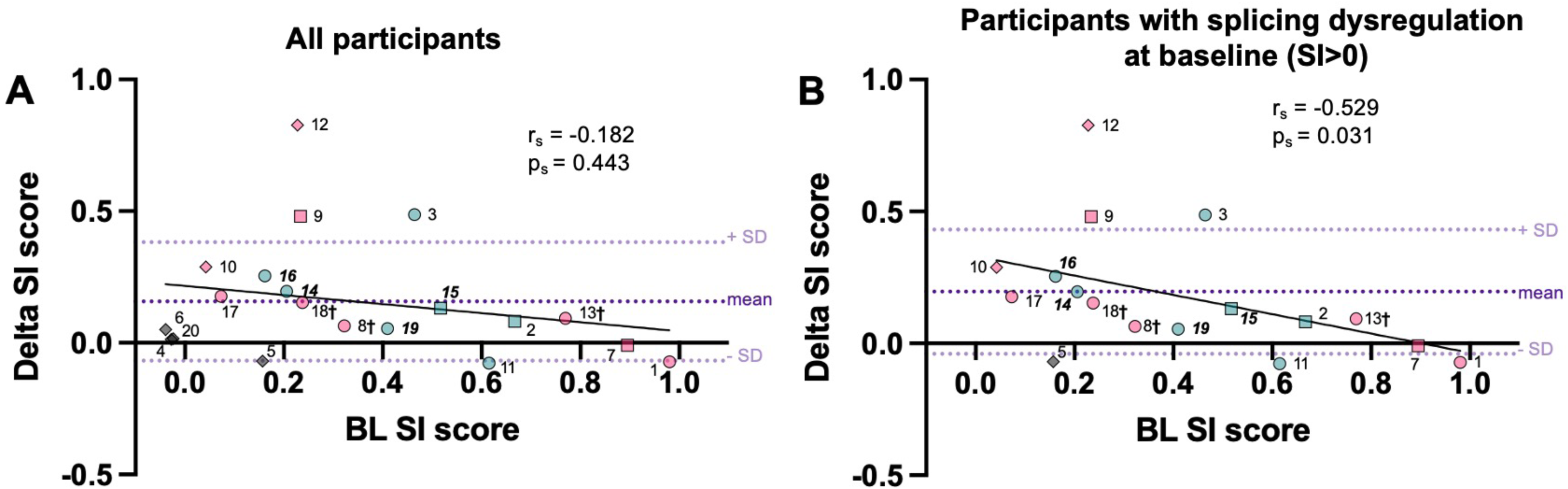
Changes in splicing index over 3 years based on baseline splicing index. **A.** Spearman correlation in all participants. Dark purple dotted line represents mean and light purple dotted lines represent ± standard deviation. Shapes represent their number of CTG repeats <300 (diamond), 300-800 (circle), >800 (square) while colors represent their phenotype juvenile (green), adult (pink), late (grey), participants with interruption of their repeats have cross next to their ID in their labels and ***bold in italic*** ID labels are participants <40 years old. **B.** Spearman correlation in all participants with splicing dysregulation at baseline (SI >0, n=17). Dark purple dotted line represents mean and light purple dotted lines represent ± standard deviation. Shapes represent their number of CTG repeats <300 (diamond), 300-800 (circle), >800 (square) while colors represent their phenotype juvenile (green), adult (pink), late (grey), participants with interruption of their repeats have cross next to their ID in their labels and ***bold in italic*** ID labels are participants <40 years old. Abbreviations: SI: splice index, BL: baseline, delta SI score: changes in SI score over 3 years, SD: standard deviation.

**Table 3:**
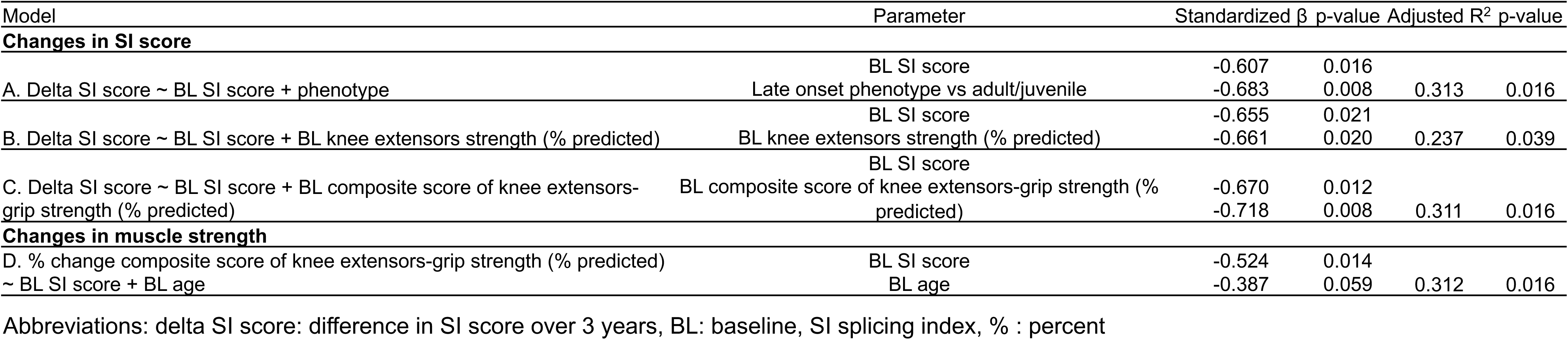
Regression model of changes in SI score and muscle strength over 3 years.

Interestingly, participants with lower strength of knee extensors muscle group (Standardized β = −0.661, p = 0.020) and SI score at baseline (Standardized β = −0.655, p = 0.021) presented with greater worsening of SI score, with these two variables explaining 24% of the variance (**Table 3 model B**). Similar results were obtained using strength values of the ankle dorsiflexors or grip strength (**Supplemental Table 1 models A-B**). However, a model using the composite score of knee extensors and grip strength explained more variance in SI score worsening over time than models including either muscle group alone (31%; **Table 3 model C vs B or Supplemental Table 1 model B**). Similar results were obtained with the other composite scores (**Supplemental Table 1 models C-E**).

In this cohort, knee extensors and grip strength significantly decreased over 3 years, whereas ankle dorsiflexors strength decreased only numerically (**Table 2**). When participants were separated by SI group (mild, moderate, and severe), changes in grip strength trended towards a decrease over three years (**Figure 2B**). Changes in strength of the composite score of the knee extensors and grip also trended in the same direction (**Supplemental Figure 3**). To verify whether the SI score at baseline is a contributing factor of changes in clinical values, multiple additional regression models were generated (**Table 3 model D** & **Supplemental Table 1 models F-G**). Older participants or those with a higher baseline SI score presented a higher decrease in the composite score of knee extensors and grip strength over time, and these variables explained 31% of the variance (**Table 3 model D**). Similar results were obtained with baseline age and change in knee extensors or grip strength over time (**Supplemental Table 1 models F-G**).

## Discussion

Despite the biomarker potential of alternative splicing and the SI in DM1, very little is known about its progression over long periods of time and across different muscle groups. Here we determined for the first time the associations between the SI score from *vastus lateralis* (VL) muscle and clinical outcomes over a 3-year period in a cohort of DM1 patients previously well characterized for clinical changes. We found that SI score in these DM1 patients worsened over 3 years overall, similar to clinical declines in strength, but with very different SI score progression from one participant to the next. Phenotype and SI score at baseline were identified as factors influencing SI progression over 3 years. This parallels previously characterized clinical progression in this population that is influenced by phenotypes(16, 17). Taken together these data support the need to integrate clinical phenotype and baseline molecular severity when interpreting longitudinal SI change.

Here, we measured SI score in the VL, a muscle with significant impact on patient daily outcomes(18, 19), and which has been relatively less studied in DM1 compared with the distal *tibialis anterior* (TA) muscle. SI score in the VL correlates with multiple clinical measures, including muscle strength, which mirrors reported SI score associations in TA muscle. As expected, our VL SI score correlates more strongly with knee extensors (proximal muscle) strength (r = −0.684) whereas TA SI score(14) correlated more strongly with ankle dorsiflexors (distal muscle) strength (r = −0.719). Regardless of individual correlations for TA or VL, the SI score in both muscles correlated strongly and negatively with muscle strength in both studies(14) with correlation coefficients values around −0.7. There was no association between SI score and 10mWT (Figure 1D). Our test protocol didn’t allow running in contrast to Provenzano *et al*(14), which allowed running and might explain why they observed a correlation and we didn’t. To better capture multisystemic involvement, we generated a composite muscle group score and compare correlations with the SI score. Interestingly we found that the composite score provides stronger correlations for both our cohort and the Provenzano *et al* cohort(14) (**Figure 1** & **Supplemental Figure 1**). Taken together these studies show that an SI score derived from either VL or TA correlates splicing and clinical outcomes, with a composite score providing stronger associations with splicing than individual measures.

Splicing dysregulation as assessed by SI score worsened over three years in our cohort, although progression was highly variable between individual participants. As expected, this increase of median SI score over 3 years at the group level from 0.24 to 0.44 was bigger than the increase reported by Provenzano *et al* over 3 months (∼0.1 mean SI change). Interestingly, participants with no splicing dysregulation at baseline (SI ≤0) do not show any significant progression over three years and were made up primarily of participants with a late-onset phenotype. Remarkably, Provenzano *et al*(14), found the biggest changes in SI score in the moderate SI score group (0.4-0.75), whereas in our study the mild SI score group (<0.4) showed the biggest change. However, in both studies there was not a change in SI score for participants in the severe SI score group, suggesting a potential ceiling effect.

Multiple regression models confirm that phenotype and baseline SI score both contribute to worsening in SI score. Clinical variables such as muscle strength at baseline also contribute to changes in SI score with a composite muscle group score explaining a larger part of the SI score variability compared to individual strength measures. Additionally, given the significant decline in grip and knee extensors strength over 3 years, we examined if baseline SI score could explain the progression of muscle strength over time. Interestingly, our regression models reveal that both SI score and age at baseline could be used to explain variability of knee extensors strength and grip individually or when used in combination in a composite score.

Our mild SI score group contains both participants with the biggest change in SI score and participants whose baseline splicing does not change significantly over time. To better understand these groups, we correlated baseline SI score with changes in SI score over 3 years. Interestingly focusing only on participants exhibiting splicing dysregulation at baseline (SI>0) greatly improved the correlation, which went from r = −0.182 to r = −0.529 (**Figure 3**). It is important to note that while there is little change in SI score over three years for these participants, two individuals (#6 & 20) did show declines in muscle strength measures during that period of time (**Figure 2 & Supplemental Figure 3**). These differences between the progression of SI score and muscle strength across different individuals highlight the complex and variable nature of pathogenesis in DM1. While limited sampling of the VL could be contributing to the variability, this issue would also similarly affect published TA muscle data.

This study identified factors such as phenotype and baseline SI score that explain changes in splicing over time, which will be important to consider during clinical trials. In looking at our cohort, participants can be divided into three groups: (1) participant whose SI score does not change significantly over time (SI lower than the mean) which is composed mostly of participants with late-onset phenotype, and participants with high SI score at baseline; (2) participants with moderate changes over 3 years (SI over the mean but lower than the mean + SD) which is composed of participants with mild SI score at baseline; and (3) participants whose SI score increased greatly over three years. The reason for this dramatic change in the latter group is currently unknown, but warrants further detailed investigation, as large fluctuation in SI scores may complicate the assessment of treatment response in clinical trials by masking therapeutic effects and creating the appearance of nonresponse. Similarly, participants in the first group with no splicing dysregulation at baseline (SI ≤ 0) will not show SI progression over time and would not be predicted to show SI score improvement following treatment. Based on our findings, baseline SI scores, composite strength measures (knee extensors, grip and ankle dorsiflexors), and clinically standardized phenotypes are important variables to be considered for future clinical trials.

### Strengths and limitations

This study is the first showing progression of splicing over 3 years in DM1 participants. Given that the sample size of our study was small, caution should be taken in the interpretation of the results until replicated with more participants. However, these replicative studies may also be limited by the difficulties in obtaining multiple longitudinal muscle biopsies, especially for a rare disease like DM1. While the fact that VL is less affected in DM1 is a limitation, both VL and TA show 3-year strength declines and VL has significant impact on patient daily outcomes(18, 19).

Furthermore, as the VL is less affected, the quality of the muscle biopsied is better, as illustrated by our ability to extract RNA and make libraries from all biopsies. Our VL findings on the associations between SI score and strength align with those previously seen in the TA, providing a choice in muscle group sampling that may be important for future studies requiring repeated muscle biopsies. Another limitation is that our observed SI changes could result from within-muscle variability rather than longitudinal changes, as the biopsy was not taken at the same site of the same muscle over time and it is also unclear if SI is uniform in a muscle. However, given the SI score changes are associated with clinical changes, we are confident in our conclusions on longitudinal changes. Also, the standard operating procedure (SOP) was the same for both biopsies, and the biopsies were performed by the same physicians.

### Conclusion

In this longitudinal study of myotonic dystrophy type 1, we observed that the SI score in the VL generally worsened over a 3-year period. While SI progression is different between participants, we identified factors in participants, such as late-onset phenotype or high baseline SI score, that contribute to reduced changes in SI score over 3 years. While these findings need to be replicated with larger cohorts, they have important implications for cohort selection for future biomarker and therapeutic clinical trials for DM1.

## Methods

### Sex as a biological variable

The study design accounted for sex as a biological variable, and both men and women were included.

### Participants

This study was part of a larger longitudinal study that began in 2002 and utilized data from the two latest collection times(16) (T3: 2015-2016 and T4: 2019). Participants with a genetic diagnosis of DM1, between 18 and 70 years old and able to provide informed consent were recruited at the neuromuscular clinic of the Santé Québec Saguenay–Lac-St-Jean – Universitaire. Participants were excluded if they had any other neuromuscular diseases or other diseases affecting function, or if they had contraindications to physical evaluation or muscle biopsy. We selected a subset of 20 participants based on the availability of their clinical data and sufficient flash-frozen tissue from VL biopsy remaining.

### Clinical assessments

Participant characteristics were either acquired during the larger longitudinal study or from clinical files. All clinical phenotypes were reviewed by the neuromuscular clinic’s neurologist. Physical evaluations and muscle biopsies were done on separate visits, with the physical evaluation done first. Maximum isometric muscle strength was measured with quantified muscle testing (QMT) with two trials on each side, unless a participant had an injury preventing the evaluation. Another trial was added if there was more than a 10% difference between the two trials (up to a maximum of six trials for each muscle group). Evaluators gave energetic standardized encouragement and contraction lasted for at least 10 seconds. Importantly, the validity and reliability of this outcome measure have been documented in DM1(20). A Jamar hand dynamometer (JLW instruments, Chicago, USA) was used to measure prehension strength and a Medup linear handheld electronic dynamometer (Atlas medic, Québec, Canada) was used for all other muscles groups following the procedure described in Hébert et al(21). Percent predicted grip strength values were computed using normative values in Mathiowetz V, *et al* (22), while other percent predicted strength values are calculated with normative values by Hébert et al (unpublished data). Maximal walking speed was measured with the 10-meter Walk Test (10mWT). Participants were asked to walk at their maximum speed without running on 14 m and could use their walking aids. The timer started at 2 m and stopped at 12 m to reduce bias due to acceleration and deceleration.

### Muscles biopsies

VL biopsies were collected with a modified Bergstrom technique by general practitioners and trained members of the research teams as previously described(15), flash frozen in liquid nitrogen and kept at −80°C until use.

### Targeted-Seq library preparation

RNA was extracted successfully from 5 mg of the 40 VL biopsies (20 participants at 2 time points) with the 2mm beads (Zymo Research) and Zymo Quick RNA Miniprep kit (Zymo Research) following manufacturer protocol including DNAse treatment. Fragment analyzer was used to assess RNA quality (Agilent). Targeted RNA sequencing analysis was done based on the previously method described in Provenzano *et al*. SuperScript® IV Reverse Transcriptase with random priming was used to make cDNA from 10 ng of RNA. Qubit (Invitrogen) and Fragment Analyzer (Agilent) was used to quantify and assessed quality of the libraries. They were equimolarly pooled and sequenced (paired reads 150bp) on an Illumina NextSeq 2000 sequencer.

### Targeted RNA-Seq analysis

The analysis was done as previously described in Provenzano *et al*. Reads were aligned to the custom reference genome found with HISAT2 v2.1.0 using the no-splice alignment and no-softclip options. Samtools v1.7, Bash CLI and a custom python script were used to get PSI values for each event. A normalized PSI value was then computed for each event using the following formula (PSI-PSI median controls)/(PSI 95^th^ percentile of DM1 – PSI median controls) with median controls and 95^th^ percentile of DM1 values pulled from Provenzano *et al*. The SI score was then computed by averaging all normalized PSI values for each participant. Note that unlike Provenzano *et al*, SI scores below 0 were not rounded up to 0 nor were SI scores over 1 rounded down to 1.

### Statistics

The Spearman rho (r_s_) correlation coefficient was used to assess the associations between SI and clinical variables. Wilcoxon signed-rank tests were used to compare baseline and 3-year values. Percent change of SI score and clinical variables was calculated using the following formula: (3-year value – baseline value)/baseline value*100. Multiple linear regression models were used to identify factors associated with the 3-year changes in SI score. Phenotype, baseline SI score and clinical variables with a Spearman correlation coefficient >0.2 were included in the models. Graph Pad Prism version 10.6.10 was used for all statistical analysis and graphs, except multiple linear regression analyses that were conducted using R 4.4.0, standardized beta coefficients were computed using the R package lm.beta and R Studio 2025.05.1.

### Study approval

The ethics review board of the Santé Québec Saguenay–Lac-St-Jean – Universitaire approved the study (2023-042) and all participants provided written informed consent.

## Data availability

SI values for each participant are available in Supplemental table 2.

## Authors contributions

CL performed statistical analysis, generated figures and wrote the manuscript. LP performed bioinformatics and statistical analysis, generated figures and revised the manuscript. RM prepared targeted libraries and revised the manuscript. JR, SC, LD and FN contributed to the study design and revised the manuscript. MPR collected clinical data and revised the manuscript. CG, JDC, JAB and ED, designed the study, analyzed the results and revised the manuscript.

## Funding support

Research in the Berglund lab is supported by funding from: the National Institutes of Health Paul D Wellstone Muscular Dystrophy Research Center at VCU & UAlbany (P50AR085906), and R01NS120485 to J.A.B. & J.D.C; the Congressionally Directed Medical Research Program (HT94252410190), the University at Albany, and the Marigold New York Myotonic Dystrophy Center fund to J.A.B & J.D.C. CL is supported by a postdoctoral fellowship from the Fonds de la recherche du Québec en santé (FRQS-#311324, #355849), AFM-Telethon (24283) and Myotonic Dystrophy Foundation (103191). ED is a holder of the Canada Research Chair (Tier 2) in Translational Science for Rare Neuromuscular Diseases. This study is also supported by funding from the Programme de soutien au développement de la mission universitaire: Enseignement et recherche du CIUSSS-SLSJ, the Marigold Foundation and Vertex Pharmaceuticals. The content is solely the responsibility of the authors and does not necessarily represent the official views of the National Institutes of Health or any other funding agency.

## Acknowledgments

We would like to thank all participants and their families. We thank all members of the Berglund and Duchesne labs as well as colleagues of the RNA institute for their helpful discussions.

## Conflict of Interest

J.A.B. serves on the Scientific Advisory Committee and Board for the Myotonic Dystrophy Foundation, has provided consultation for D. E. Shaw Research, Dyne Therapeutics, Entrada Therapeutics, Juvena Therapeutics, Kate Therapeutics, Syros Pharmaceuticals, Sanofi, and Wayfinder Biosciences, and received research funding from Agios Pharmaceuticals, Arthex, Biomarin Pharmaceuticals, PepGen, Syros Pharmaceuticals, and Vertex Pharmaceuticals. J.D.C. is a part-time employee of the University of Florida’s Center for NeuroGenetics and J.A.B. receives licensing royalties from the University of Florida. J.D.C and J.A.B. have intellectual property filings related to DM and DM therapeutics and equity in Repeat RNA Therapeutics. ED consulted or currently consults for Dyne Therapeutics, PepGen, Avidity, AMO Pharma and Vertex Pharmaceuticals, and received research funding from Vertex Pharmaceuticals. JR, SC, LMD, and FN are employees of Vertex Pharmaceuticals and hold stock and/or stock options in the company.

