## Supplemental Figure 1 for "Three-Year Longitudinal Analysis of Splicing Changes in Myotonic Dystrophy Type 1"

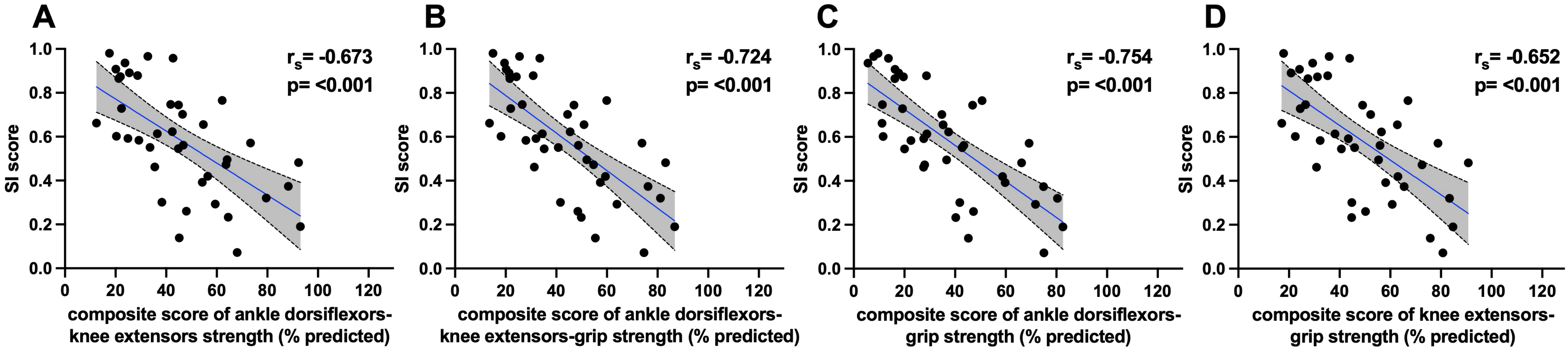


**Supplemental Figure 1: Spearman correlations of splicing index and composite scores in Provenzano et al dataset to validate the approach. A.** Correlation of splicing index and composite score average of percent predicted quantitative muscle testing of ankle dorsiflexors and knee extensors strength in Provenzano *et al* dataset. **B.** Correlation of splicing index and composite score average of percent predicted quantitative muscle testing of ankle dorsiflexors, knee extensors and grip strength in Provenzano *et al* dataset. **C.** Correlation of splicing index and composite score average of percent predicted quantitative muscle testing of ankle dorsiflexors and grip strength in Provenzano *et al* dataset. **D.** Correlation of splicing index and composite score average of percent predicted quantitative muscle testing of knee extensors and grip strength in Provenzano *et al* dataset. Abbreviations: BL: baseline, SI: splicing index, %: percent
