## Supplemental Figure 2 for "Three-Year Longitudinal Analysis of Splicing Changes in Myotonic Dystrophy Type 1"

| | Delta SI score | % change ankle dorsi | % change knee ext | % change pinch | % change grip | % change composite (ankle dorsi-knee ext) | % change composite (ankle dorsi-knee ext-grip) | % change composite (ankle dorsi-grip) | %change composite (knee ext-grip) |
|---|---|---|---|---|---|---|---|---|---|
| **BL age** | -0.30 | 0.03 | -0.20 | -0.27 | -0.23 | -0.08 | -0.11 | 0.04 | -0.20 |
| **BL CTG** | 0.17 | 0.04 | -0.12 | 0.14 | -0.12 | -0.02 | -0.10 | -0.13 | -0.25 |
| **BL SI score** | -0.18 | 0.17 | -0.34 | 0.16 | -0.42 | -0.06 | -0.25 | -0.12 | -0.51* |
| **BL knee Ext PP** | -0.22 | 0.12 | 0.13 | -0.01 | 0.19 | 0.16 | 0.25 | 0.23 | 0.34 |
| **BL ankle PP** | -0.28 | 0.24 | 0.50* | 0.45 | 0.34 | 0.26 | 0.39 | 0.40 | 0.35 |
| **BL grip PP** | -0.38 | 0.22 | 0.52* | -0.04 | 0.05 | 0.18 | 0.19 | 0.24 | 0.20 |
| **BL composite (ankle dorsi-knee ext)** | -0.26 | 0.19 | 0.34 | 0.27 | 0.36 | 0.26 | 0.39 | 0.35 | 0.43 |
| **BL composite (ankle dorsi-knee ext-grip)** | -0.35 | 0.36 | 0.45 | 0.25 | 0.21 | 0.33 | 0.41 | 0.43 | 0.34 |
| **BL composite (ankle dorsi-grip)** | -0.31 | 0.30 | 0.54* | 0.23 | 0.16 | 0.25 | 0.32 | 0.36 | 0.24 |
| **BL composite (knee ext-grip)** | -0.28 | 0.25 | 0.42 | -0.06 | 0.18 | 0.23 | 0.29 | 0.30 | 0.36 |

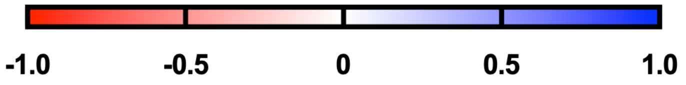


**Supplemental figure 2: Spearman correlations of splicing index and clinical values at baseline and changes over 3 years.** *$p<0.05$, abbreviations: BL: baseline, CTG: number of CTG repeats, SI: splicing index, % change ankle dorsi: percent change over 3 years of ankle dorsiflexors strength, % change knee ext: percent change over 3 years of knee extensors strength, % change pinch: percent change over 3 years of pinch strength, % change grip: percent change over 3 years of grip strength, % change composite (ankle dorsi-knee ext): percent change over 3 years of composite score average of percent predicted quantitative muscle testing of ankle dorsiflexors and knee extensors, % change composite (ankle dorsi-knee ext-grip): percent change over 3 years of composite score average of percent predicted quantitative muscle testing of ankle dorsiflexors, knee extensors and grip, % change composite (ankle dorsi-grip): percent change over 3 years of composite score average of percent predicted quantitative muscle testing of ankle dorsiflexors and grip, % change composite (knee ext-grip): percent change over 3 years of composite score average of percent predicted quantitative muscle testing of knee extensors and grip, knee ext PP: knee extensors percent predicted quantitative muscle testing strength, ankle PP: ankle dorsiflexors percent predicted quantitative muscle testing strength, grip pp: grip percent predicted quantitative muscle testing strength, composite (ankle dorsi-knee ext): composite score average of percent predicted quantitative muscle testing of ankle dorsiflexors and knee extensors strength, composite (ankle dorsi-knee ext-grip): composite score average of percent predicted quantitative muscle testing of ankle dorsiflexors, knee extensors and grip strength, composite (ankle dorsi-grip): composite score average of percent predicted quantitative muscle testing of ankle dorsiflexors and grip strength, composite (knee ext-grip): composite score average of percent predicted quantitative muscle testing of knee extensors and grip strength.
