## Supplemental Figure 3 for "Three-Year Longitudinal Analysis of Splicing Changes in Myotonic Dystrophy Type 1"

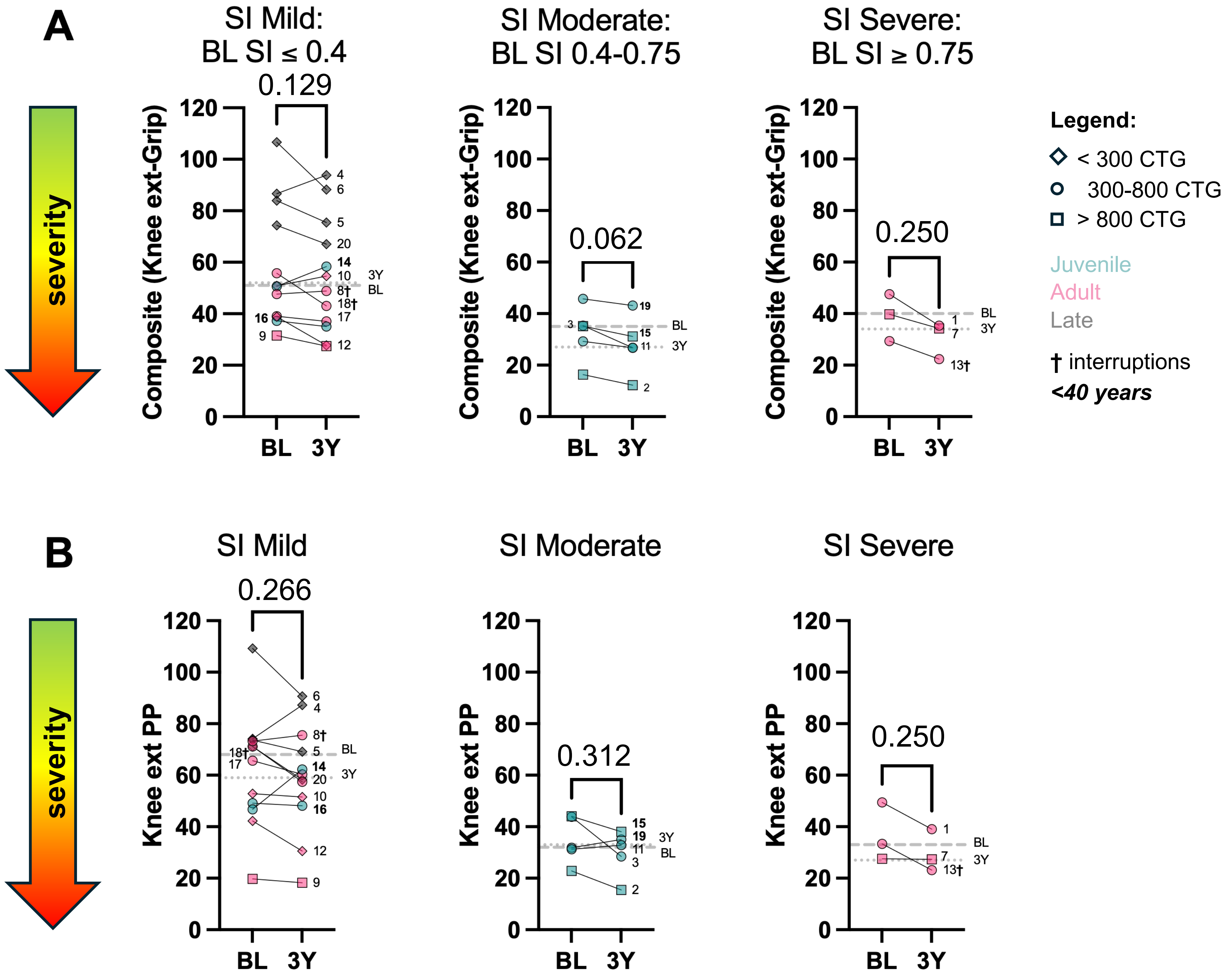


**Supplemental Figure 3: composite score of knee extensors and grip strength and knee extensors strength changes over 3 years. A.** Composite score of knee extensors and grip strength changes over 3 years. Shapes represent their CTG repeats <300 (diamond), 300-800 (circle), >800 (square) while colors represent their phenotype juvenile (green), adult (pink), late (grey), participants with interruption of their repeats have cross next to their ID in their labels and ***bold in italic*** ID labels are participants <40 years old. Median are presented as grey lines large dash= baseline, smaller dash=3 years. . **B.** Changes in knee extensors strength over 3 years. Shapes represent their CTG repeats <300 (diamond), 300-800 (circle), >800 (square) while colors represent their phenotype juvenile (green), adult (pink), late (grey), participants with interruption of their repeats have cross next to their ID in their labels and ***bold in italic*** ID labels are participants <40 years old. Median are presented as grey lines large dash= baseline, smaller dash=3 years. Abbreviations: SI: splice index, BL: baseline, 3Y: 3 years, composite (knee ext-grip): composite score average of percent predicted quantitative muscle testing of knee extensors and grip strength, knee ext pp: knee extensors percent predicted quantitative muscle testing strength, CTG: number of CTG repeats. *p<0.05 Wilcoxon test.
