## Supplemental Table 1 for "Three-Year Longitudinal Analysis of Splicing Changes in Myotonic Dystrophy Type 1"

Supplemental Table 1: Additional regression models of changes in SI score and muscle strength over 3 years

| Model | Parameter | Standardized β | p-value | Adjusted $R^2$ | p-value |
|---|---|---|---|---|---|
| **Changes in SI score** | | | | | |
| A. Delta SI score ~ BL SI score + BL ankle dorsiflexors strength (% predicted) | BL SI score | -0.510 | 0.047 | | |
| | BL ankle dorsiflexors strength (% predicted) | -0.585 | 0.026 | 0.226 | 0.050 |
| B. Delta SI score ~ BL SI score + BL grip strength (% predicted) | BL SI score | -0.482 | 0.053 | | |
| | BL grip strength (% predicted) | -0.530 | 0.036 | 0.191 | 0.064 |
| C. Delta SI score ~ BL SI score + BL composite score of ankle dorsiflexors-knee extensors strength (% predicted) | BL SI score | -0.683 | 0.012 | | |
| | BL composite score of ankle dorsiflexors-knee extensors strength (% predicted) | -0.757 | 0.006 | 0.341 | 0.014 |
| D. Delta SI score ~ BL SI score + BL composite score of ankle dorsiflexors-knee extensors-grip strength (% predicted) | BL SI score | -0.628 | 0.016 | | |
| | BL composite score of ankle dorsiflexors-knee extensors-grip strength (% predicted) | -0.718 | 0.007 | 0.330 | 0.016 |
| E. Delta SI score ~ BL SI score + BL composite score of ankle dorsiflexors-grip strength (% predicted) | BL SI score | -0.501 | 0.047 | | |
| | BL composite score of ankle dorsiflexors-grip strength (% predicted) | -0.591 | 0.022 | 0.239 | 0.044 |
| **Changes in muscle strength** | | | | | |
| F. % change knee extensors strength ~ BL SI score + BL age | BL SI score | -0.320 | 0.136 | | |
| | BL age | -0.476 | 0.032 | 0.216 | 0.049 |
| G. % change grip strength ~ BL SI score + BL age | BL SI score | -0.440 | 0.046 | | |
| | BL age | -0.365 | 0.092 | 0.212 | 0.051 |

Abbreviations: delta SI score: difference in SI score over 3 years, BL: baseline, SI splicing index, % : percent
