## Supplemental Table 2 for "Three-Year Longitudinal Analysis of Splicing Changes in Myotonic Dystrophy Type 1"

Supplemental Table 2: SI scores

| ID | TA normalized BL SI | TA normalized 3Y SI | Changes in TA normalized SI |
|---|---|---|---|
| 1 | 0.980 | 0.909 | -0.072 |
| 7 | 0.895 | 0.884 | -0.01 |
| 13 | 0.770 | 0.863 | 0.093 |
| 2 | 0.667 | 0.749 | 0.082 |
| 11 | 0.615 | 0.538 | -0.077 |
| 15 | 0.517 | 0.649 | 0.132 |
| 3 | 0.464 | 0.951 | 0.487 |
| 19 | 0.41 | 0.464 | 0.054 |
| 8 | 0.322 | 0.386 | 0.064 |
| 18 | 0.238 | 0.391 | 0.153 |
| 9 | 0.234 | 0.714 | 0.480 |
| 12 | 0.228 | 1.056 | 0.827 |
| 14 | 0.206 | 0.402 | 0.196 |
| 16 | 0.162 | 0.416 | 0.254 |
| 5 | 0.157 | 0.087 | -0.07 |
| 17 | 0.073 | 0.249 | 0.177 |
| 10 | 0.042 | 0.33 | 0.288 |
| 20 | -0.024 | -0.008 | 0.016 |
| 4 | -0.028 | -0.016 | 0.013 |
| 6 | -0.039 | 0.011 | 0.05 |
| Median | 0.236 | 0.440 | 0.088 |
| Q1, Q3 | 0.094, 0.591 | 0.269, 0.835 | 0.013, 0.240 |

Abbreviations: TA: *tibialis anterior*, BL: baseline, SI: splicing index, 3Y: 3 years
